# VIRULIGN: fast codon-correct alignment and annotation of viral genomes

**DOI:** 10.1101/409052

**Authors:** Pieter Libin, Koen Deforche, Ana B. Abecasis, Kristof Theys

## Abstract

Virus sequence data are an essential resource for reconstructing spatiotemporal dynamics of viral spread as well as to inform treatment and prevention strategies. However, the potential benefit for these applications critically depends on accurate and correctly annotated alignments of genetically heterogeneous data. VIRULIGN was built for fast codon-correct alignments of large datasets, with standardized genome annotation and various alignment export formats.

VIRULIGN is freely available at https://github.com/rega-cev/virulign as an open source software project.

## 1 Introduction

Many viral pathogens, in particular RNA viruses, are fast evolving and exhibit extensive genetic diversity within and between hosts. High rates of evolution provide a mechanism for rapid adaptation to changing environmental conditions and escape from selective pressures. Structural, functional and phenotypic predictions based on the viral genotype have important applications in drug design, diagnostics and clinical management of viral infections. Moreover, virus genetic data are a requisite for phylogenetic inference of past evolutionary events and active epidemiological surveillance.

These genotype-based applications strongly depend on accurate sequence alignments. Additionally, protein coding sequences (CDS) should be analyzed in the correct open reading frame (ORF) to preserve the underlying biological relevance. Multiple sequence alignments (MSA) of viral pathogens are typically constructed by progressive-iterative approaches such as MAFFT, MUSCLE or Clustal Omega [8,6,19]. However, these heuristic methods are less capable of mitigating camshaft errors and can be sensitive to noise in the sequence data. Alternatively, guidance of the alignment process by a reference sequence can overcome these limitations [25]. Here we present VIRULIGN, a fast codon-correct alignment and genome annotation tool that can be applied to sequence data from different viral pathogens.

## 2 Features

VIRULIGN is a cross-platform (GNU/Linux, Unix, MacOS and Windows) and easy-to-use command line application that can handle large sequence datasets in a computationally efficient manner. VIRULIGN aligns sequences and corrects the alignment for codon anomalies resulting from single nucleotide alterations. Automated frame shift correction and genome annotation increases the quality of the alignment and reduces the need for manual editing, thereby addressing the need for reproducible research [16].

A codon-correct MSA is essential for evolutionary hypothesis testing and phylogenetic inference using codon substitution models [18] and for detecting footprints of selective constraints on coding sequence alignments. In addition, the identification of amino acid mutations (including insertions or deletions) associated with drug resistance (HIV-1, Hepatitis C virus, Influenza virus), disease outcome (Hepatitis B virus) or epidemic potential (Ebola virus, Chikungunya virus) are important aspects in the management of infectious diseases.

VIRULIGN offers standardized protein annotation of the target CDS, relative to positions within a curated reference genome, through the use of an XML file. This annotation file can be easily defined by the user and VIRULIGN provides pre-defined annotations for several viral pathogens (see Supplementary Information, SI). This feature facilitates genome-wide or protein-specific analyses and enables to optimize reference sequences in terms of completeness and representativeness [23].

VIRULIGN provides support to export the computed alignment to various output formats, where different options can be combined to obtain an appropriate alignment representation. Alignments can be exported, either in nucleotide or amino acid alphabet, as fasta and CSV files, with the latter representing protein positions and mutations as separate columns.

VIRULIGN is an open-source project (GPLv2 license) written in the C++ programming language. VIRULIGN was previously used in different research areas in infectious diseases (see SI), and can be easily integrated in data management and analysis platforms for viral pathogens.

## 3 Methods

VIRULIGN attempts to construct a MSA of a set of target sequences 𝒯 with respect to a reference sequence *r* (Figure of the alignment process in SI). For each target sequence *t* ∈ 𝒯, a codon correct pairwise alignment with *r* is computed. During this procedure, different alignments are performed using the Needleman-Wunsch global alignment algorithm [13]. We will refer to the amino acid representation of the reference sequence *r* as *AA*(*r*).

Firstly, we perform a Needleman-Wunsch nucleotide alignment of *r* and *t*, resulting in alignment 𝒜_*nt*_(*r, t*). Secondly, the three ORFs of target sequence *t* are translated to their respective amino acid representation. Each of these amino acid sequences is aligned to *AA*(*r*) using the Needleman-Wunsch algorithm. From these three alignments, the alignment with the highest alignment score is selected. This amino acid alignment is than converted to a nucleotide alignment 𝒜_*cc*_(*r, t*), by replacing each of the amino acids with their respective nucleotide codon. Thirdly, if 𝒜_*cc*_(*r, t*) and 𝒜_*nt*_(*r, t*) differ, we suspect that a frame-shift has occurred in the target sequence. We then attempt to fix the frame-shift, by detecting the first isolated gap of which the size is not a multiple of three, and replace it by an ‘n’ nucleotide symbol. Finally we move again to the second step and the procedure is repeated until no more frame-shifts are detected, or the maximum number of frame-shifts (i.e., a configuration option) has been exceeded. In the latter case, target sequence *t* is excluded from the MSA.

This procedure results in a set of codon-correct aligned target sequences, where each of these alignments contain information about possible insertions in the target sequence. This data structure can be exported to a MSA, in a variety of output formats (see Features section), by iterating over the alignment columns in each of the pairwise alignments.

We derive VIRULIGN’s computational complexity in the SI.

## 4 Application and future perspectives

We demonstrate VIRULIGN by constructing MSAs of data collected from public databases for HIV-1, Dengue virus (DENV) and Zika virus (ZIKV). Detailed information on the datasets and methods used is available as SI.

Firstly, full-length genomes from different genotypes of DENV serotype 1 (n=1433), the most common mosquito-borne viral infection globally, were aligned with VIRULIGN, MAFFT, MUSCLE and Clustal Omega. This example shows that, compared to the other tools, VIRULIGN generated an amino acid alignment in the correct ORF without the need for manual correction, while remaining computationally efficient. Secondly, a selected subset of full-length ZIKV genomes (n=19) was aligned with VIRULIGN using a XML annotation file. The alignment was exported in an amino acid representation to illustrate, in conjunction with other command line utilities, the variability at a glycosylation motif that instigated the effort to correct the ZIKV reference sequence. Thirdly, we conducted two experiments in the context of HIV-1. HIV-1 exhibits three ORFs that together translate the complete set of viral proteins, however, these different reading frames complicate the alignment of the respective CDS. A curated set of full-length HIV-1 genomes (n=2966), from different sub-types, was aligned with VIRULIGN to select the gag poly-protein and identify encoded proteins in an efficient manner. Similar operations can be easily applied to other HIV-1 poly-proteins. As a second example, we used VIRULIGN to align a large HIV-1 dataset (n=111222) spanning the reverse transcriptase enzyme, for which an accurate alignment has significant clinical importance in the context of drug resistance detection. Due to its favorable computational complexity in this context, VIRULIGN performed the alignment much faster than MAFTT.

Future developments include a community-driven repository of standardized and curated genome annotations of representative reference sequences and the integration of VIRULIGN in tools for surveillance and genomic epidemiology.

## Acknowledgments

PL is funded by a doctoral grant of the Research Foundation - Flanders (FWO). We thank A.-M. Vandamme, S. Imbrechts, L. Cuypers and F. Ferreira for testing the software.

## Supplementary Information

### S1 Introduction

Virus sequence data are an essential resource for reconstructing spatiotemporal dynamics of viral spread as well as treatment and prevention strategies. However, the potential benefit of using sequence data for these applications critically depends on the accuracy and the correct annotation of these alignments of genetically diverse data. In particular, coding sequences of viral pathogens should be analyzed in their corresponding open reading frame (ORF) to fully utilize their biological information. Therefore, while the construction of multiple sequence alignments (MSAs) can be done with a range of sequence alignment softwares, MSAs of virus coding sequences in the correct reading frame and annotated with respect to the proteins encoded in the genome are more difficult to achieve.

VIRULIGN is an easy-to-use command line application to construct codon-correct alignments of large virus sequence datasets. Additionally, VIRULIGN has support for standardized genome annotation and implements various alignment export formats that are useful for various research applications. VIRULIGN is an open-source project written in the C++ programming language and available under the GPLv2 license.

VIRULIGN operates by aligning each target sequence (i.e., *t* ∈ *T*) of the input file codon-correctly against the reference sequence (*r*). Subsequently a multiple sequence alignment *MSA*(*r, T*) is constructed based on all codon-correct (cc) pairwise aligned target sequences *A*_*cc*_(*r, t*) (Figure S1).

**Figure S1:**
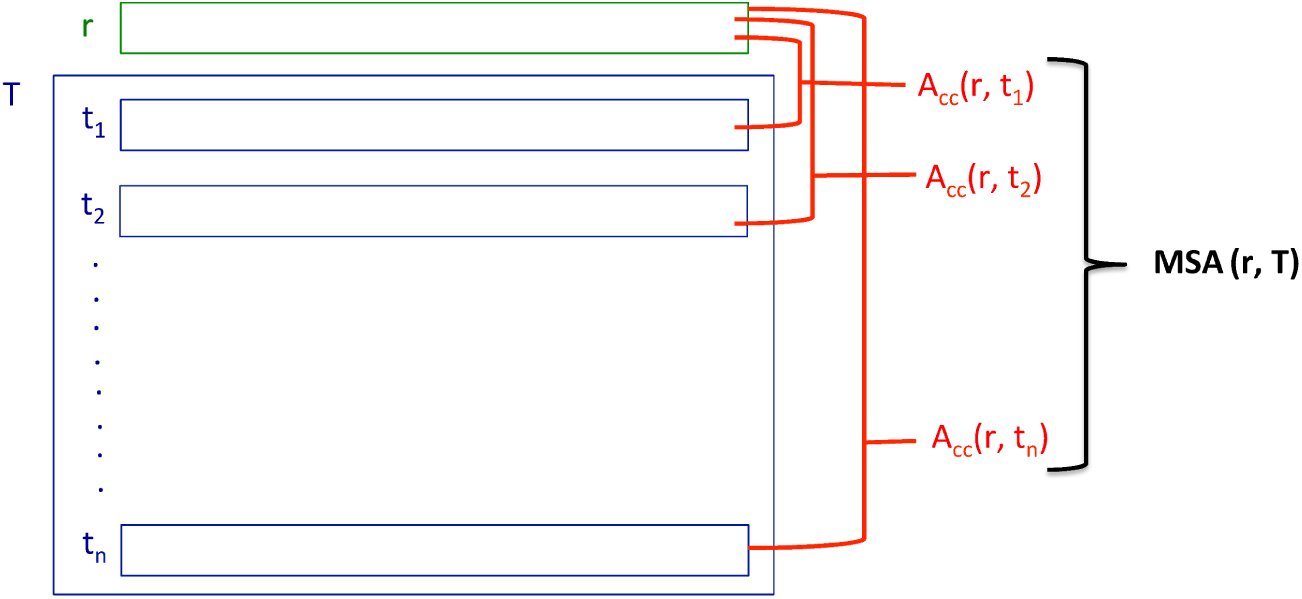
Schema of the VIRULIGN alignment process.

### S2 Installation

The most recent version and executable of VIRULIGN can be downloaded from the GitHub project web page:

https://github.com/rega-cev/virulign

Instructions are provided to build and install the software. VIRULIGN is a cross-platform application and has been tested on GNU/Linux, MacOS and Windows.

### S3 Use and features

#### Basic command

VIRULIGN minimally requires a FASTA file with target sequences and a reference sequence in order to generate a codon-correct alignment in a predefined output format (see below). The reference sequence can be either provided in FASTA format or embedded in an XML file (see below). The default command for VIRULIGN is as follows:

~~~
$ virulign [reference.fasta orf-description.xml] sequences.fasta
~~~

#### Optional arguments

Additional parameters can be specified to configure the alignment construction and to export the alignment to a variety of output formats. In case that a parameter has not been explicitly specified, the first value of this optional parameter is used as the default value. The following parameters can be used to configure the alignment and its representation ^1^:

~~~
$ virulign [reference.fasta orf-description.xml]
           sequences.fasta
    --exportKind [Mutations PairwiseAlignments
                  GlobalAlignment PositionTable
                  MutationTable]
    --exportAlphabet [AminoAcids Nucleotides]
    --exportWithInsertions [yes no]
    --exportReferenceSequence [no yes]
    --gapExtensionPenalty doubleValue=>3.3
    --gapOpenPenalty doubleValue=>10.0
    --maxFrameShifts intValue=>3
~~~

#### S3.1 Description of the optional parameters

##### S3.1.1 exportKind

The parameter `--exportKind` defines the output type of the alignment, either a FASTA sequence file or a CSV mutation file. To display the different options, consider this example FASTA input file including sequences of different length and a full-length reference sequence for comparison.

~~~
>Ref
CCCATTAGCCCTATTGAGACTGTACCAGTAAAATTAAAGCCAGGAATGGATGGCCCAAAA
>Seq1
ATTGACACTGTACCAGTAACATTAAAGCCAGGAATGGATGGACCAAAG
>Seq2
CCTATGGAAACTGTGCCAGTAAAATTAAAGCCAGGAATGGAT
>Seq3
CTCATTAGTCCTATTAGTGTAAAATTAAAACCAGGAATGGATGGCCCAAGG
>Seq4
AGTCCCATTGAAACTGTACCAGTAAAAGGAGATGGCCCAAAG
~~~

The option `GlobalAlignment` will generate a FASTA file of the target sequences aligned against a single reference sequence and formatted as a MSA.

~~~
>Ref
CCCATTAGCCCTATTGAGACTGTACCAGTAAAATTAAAGCCAGGAATGGATGGCCCAAAA
>Seq1
------------ATTGACACTGTACCAGTAACATTAAAGCCAGGAATGGATGGACCAAAG
>Seq2
CCTATG---------GAAACTGTGCCAGTAAAATTAAAGCCAGGAATGGAT---------
>Seq3
CTCATTAGTCCTATTAGT GTAAAATTAAAACCAGGAATGGATGGCCCAAGG
>Seq4
------AGTCCCATTGAAACTGTACCAGTAAAA---------GGA GATGGCCCAAAG
~~~

The option `PairwiseAlignments` will generate a FASTA file of the target sequences, with each sequence aligned separately against the reference sequence.

~~~
>Ref
CCCATTAGCCCTATTGAGACTGTACCAGTAAAATTAAAGCCAGGAATGGATGGCCCAAAA
>Seq1
------------ATTGACACTGTACCAGTAACATTAAAGCCAGGAATGGATGGACCAAAG
>Ref
CCCATTAGCCCTATTGAGACTGTACCAGTAAAATTAAAGCCAGGAATGGATGGCCCAAAA
>Seq2
CCTATG---------GAAACTGTGCCAGTAAAATTAAAGCCAGGAATGGAT---------
~~~

The option `PositionTable` will create a comma-separated value (CSV) file where each position of the alignment is given as a separate column. The CSV file is annotated according to the numerical positions in the protein (e.g., Table 1).

**Table 1:**
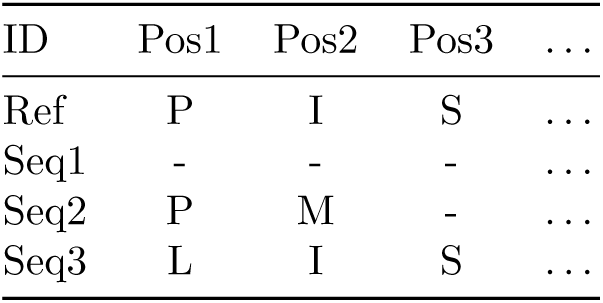
Position3Table example.

| ID | Pos1 | Pos2 | Pos3 | ... |
| --- | --- | --- | --- | --- |
| Ref | P | I | S | ... |
| Seq1 | - | - | - | ... |
| Seq2 | P | M | - | ... |
| Seq3 | L | I | S | ... |

The option `MutationTable` will create a CSV file where each mutation present at a specific position is given as a separate column in Boolean representation. The CSV file is annotated according to the numerical position in the protein (e.g., Table 2).

**Table 2:** MutationTabel example.

| ID | Mut1P | Mut1L | Mut2I | Mut2M | ... |
| --- | --- | --- | --- | --- | --- |
| Ref | y | n | y | n | ... |
| Seq1 | n | n | n | n | ... |
| Seq2 | y | y | n | y | ... |
| Seq3 | n | y | y | n | ... |

- the option `Mutations` will output, for each sequence, a list of amino acids changes compared to the reference sequence (e.g., Table 3).

**Table 3:** Mutations example.

| ID | Mutations |
| --- | --- |
| Seq1 |  |
| Seq2 | 2M |
| Seq3 | 1L |

##### S3.1.2 exportAlphabet

The parameter `--exportAlphabet` defines the alphabet in which the alignment is generated.

- the option `AminoAcids` will export an alignment with the translation of the nucleotide codons.
- the option `Nucleotides` will export an alignment with nucleotides

##### S3.1.3 exportWithInsertions

The parameter `--exportWithInsertions` determines whether insertions can be added to the reference sequence.

- the option `Yes` will insert gaps into the reference sequence to accommodate the identification of codon insertions in the target sequences.
- the option `No` removes codon insertions in the reference sequence that were generated during the alignment procedure.

##### S3.1.4 exportReferenceSequence

The parameter `--exportReferenceSequence`, controls whether the reference sequence is to be added to the alignment (Yes/No).

##### S3.1.5 gapExtensionPenalty

The parameter `--gapExtensionPenalty` defines the value of the penalty to extend an existing gap.

##### S3.1.6 gapOpenPenalty

The parameter `--gapOpenPenalty` defines the value of the penalty to start a new gap.

##### S3.1.7 maxFrameShifts

The parameter `--maxFrameShifts` defines the maximum number of frame-shifts allowed.

#### S3.2 Alignment annotation

A reference sequence can be either provided to VIRULIGN in FASTA format or embedded in an XML file. In this XML file, also an annotation of the different proteins, regions or other structures can be given by the positions relative to the reference genome.

For illustrational purposes, we present a toy XML file to align virus sequence data against a reference sequence and the annotation of the proteins A, B and C in the genome of VIRUS. Later in this document, we will consider and use some more realistic annotations (i.e., ZIKV, HIV-1).

~~~
<?xml version=“1.0” encoding=“UTF-8”?>
    <orf name=“VIRUS”
         referenceSequence=“atgaaaaacccaaaaaagaaatccgga” >
       <protein abbreviation=“A”
                startPosition=“1” stopPosition=“7” />
       <protein abbreviation=“B”
                startPosition=“7” stopPosition=“13” />
       <protein abbreviation=“C”
                startPosition=“13” stopPosition=“17” />
    </orf>
~~~

Table 4 shows an example of an alignment that is constructed with this XML file as reference, and is exported using the tabular format.

**Table 4:** CSV alignment using the toy reference XML.

| ID | A_1 | A_2 | ... | B_1 | ... |
| --- | --- | --- | --- | --- | --- |
| Seq1 | X | Y | ... | C | ... |
| Seq2 | X | T | ... | F | ... |
| Seq3 | X | Y | ... | D | ... |
| Seq4 | R | M | ... | D | ... |

### S4 Examples

We demonstrate the use of VIRULIGN by constructing sequence alignments for three viral pathogens that are the causative agents for major epidemics: HIV-1, Dengue virus serotype 1 (DENV-1) and Zika virus (ZIKV). For each pathogen, a different feature of VIRULIGN is demonstrated.

Virus sequence datasets were collected from public databases (i.e., Genbank and the Stanford HIV Drug Resistance Database) and passed to different alignment software applications for evaluation. We used a minimum number of processing steps as possible between dataset retrieval and alignment input to clearly illustrate the strength of VIRULIGN. All data files of these examples can be found on the tutorial web page:

https://github.com/rega-cev/virulign-tutorial

#### S4.1 DENV-1 alignment

We compare the output of VIRULIGN against three popular alignment tools (MAFFT, MUSCLE and Clustal Omega) in their ability to construct an accurate codon-correct alignment from a genome sequence dataset.

##### S4.1.1 Sequence dataset

Genome sequence data of DENV-1 (i.e., Dengue Serotype 1) were collected from the Dengue Virus Variation Database^2^ [7]. Only full-length nucleotide sequences originating from a human host were retained and identical sequences were collapsed. From a total of 3539 genome sequences, the corresponding serotype information was used to select a subset of 1432 DENV-1 isolates.

The input FASTA file ‘denv-1.fasta’ can be found in the tutorial Dengue folder:

https://github.com/rega-cev/virulign-tutorial/examples-alignments/DENV/

##### S4.1.2 Reference sequence

The NCBI Reference Sequence for DENV-1 is NC_001477^3^. Find a FASTA file ‘NC_001477.fasta’ that contains this reference sequence in tutorial Dengue folder:

https://github.com/rega-cev/virulign-tutorial/examples-alignments/DENV/

##### S4.1.3 Alignments by different tools

Alignments were constructed using the default parameters for each tool. The following versions were downloaded; VIRULIGN (v1.0), MAFFT (v7.313) [9], MUSCLE (v3.8.31) [6] and Clustal Omega (v1.2.3) [19].

The genome sequence of NC_001477 was added to the target dataset to facilitate the trimming of constructed alignments to the boundaries of the reference CDS, in order to remove the alignment of the 5’/3’ untranslated regions. The codon-correctness of the alignment was then evaluated by visually inspecting the amino acid translation of the respective alignments.

The following commands were used to obtain alignments:

~~~
$ mafft --auto denv-1.fasta > denv-1-mafft.fasta

$ muscle -in denv-1.fasta -out denv-1-muscle.fasta

$ clustalo --auto -i denv-1.fasta -o denv-1-clustalo.fasta

$ virulign NC_001477.fasta denv-1.fasta
    --exportKind GlobalAlignment
    --exportAlphabet Nucleotides > denv-1-virulign.fasta
~~~

Each alignment was then trimmed to the start and stop position of the coding sequence of the reference sequence, the size of this coding region is 10188 nucleotides. All trimmed alignments can be found in tutorial Dengue folder:

https://github.com/rega-cev/virulign-tutorial/examples-alignments/DENV

From this evaluation, it can be observed that VIRULIGN is able to handle insertions and deletions without disrupting the reading frame and resulting in the absence of stop codons within the alignment, while maintaining quality of the alignment. Figure S2 visualises a selected window from constructed alignments to illustrate the codon-correctness of VIRULIGN. We recorded the time needed for each alignment construction (Table 5) ^4^. This evaluation shows that VIRULIGN is able to obtain these results while still being computationally competitive with MAFFT.

**Table 5:** Run time comparison between VIRULIGN, mafft, muscle and clustalo.

| Command | Alignment | Run time |
| --- | --- | --- |
| mafft | denv-1-mafft.fasta | 2m33s |
| muscle | denv-1-muscle.fasta | 287m |
| clustalo | denv-1-clustalo.fasta | 760m |
| virulign | denv-1-virulign.fasta | 19m46s |

**Figure S2:**
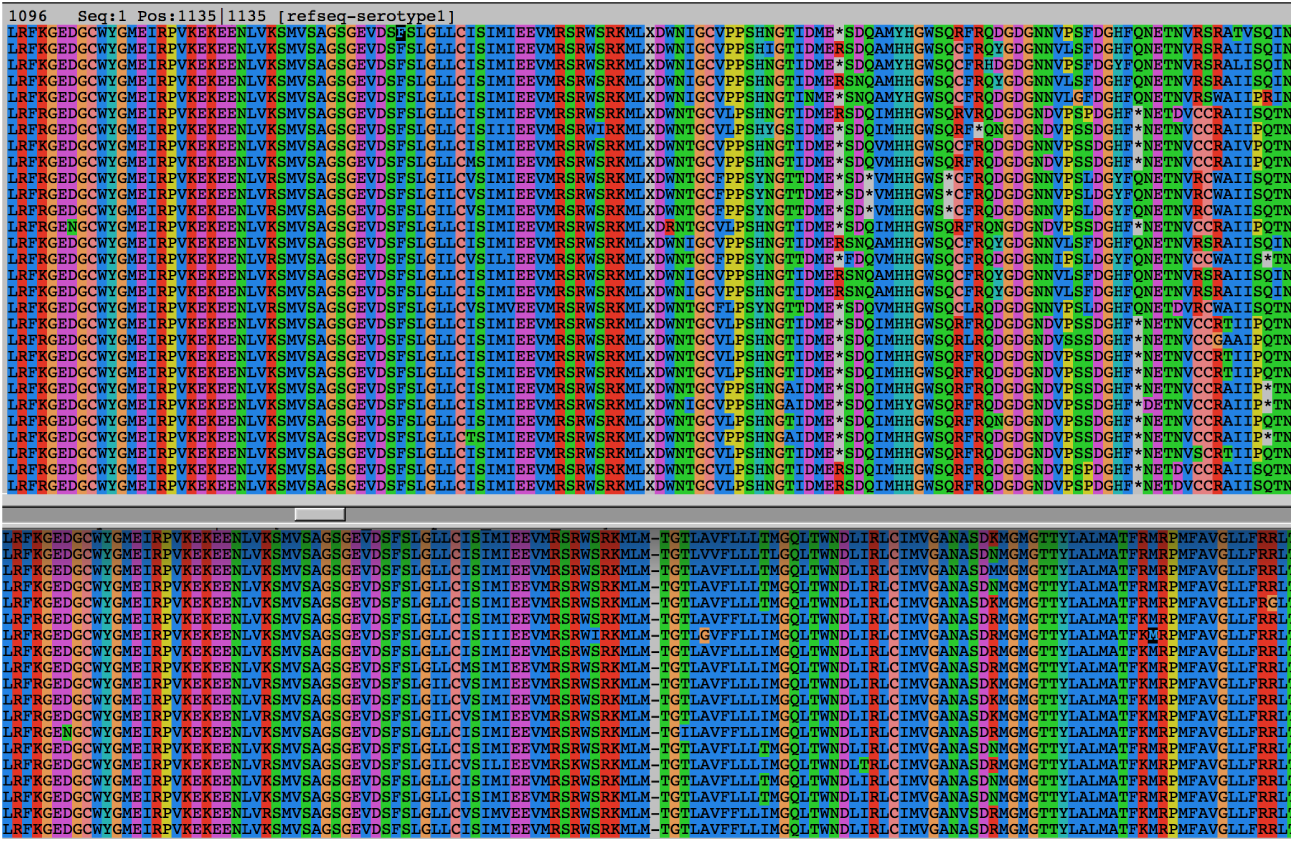
MAFFT (top window) and VIRULIGN (bottom window) constructed alignments visualized with SeaView. Stop codons are denoted by the presence of an asterisk.

##### S4.1.4 Annotation file

While the reference sequence for DENV-1 was provided by means of a simple FASTA file, VIRULIGN can also be used with an XML file containing both the reference sequence and the protein annotation of the reference genome. To construct the same alignment as above, with an annotated reference sequence, use this command:

~~~
$ virulign DENV1-NC001477.xml denv-1.fasta
    --exportKind GlobalAlignment
    --exportAlphabet Nucleotides > denv-1-virulign.fasta
~~~

XML annotation files for each of the four DENV serotypes are available in the VIRULIGN references folder:

https://github.com/rega-cev/virulign/references

More information on this XML feature is available in the section S3 Use and Features. In the next section, we demonstrate how this XML annotation file can be used to obtain alignments directed towards specific research applications.

#### S4.2 ZIKV alignment

In 2015, ZIKV caused a worldwide public health emergency, resulting in an intensive community effort to identify genomic correlates of disease manifestations of microcephaly and other neurological complications. As we have shown recently [23], the rapid advance in ZIKV genomics resulted in inconsistencies that complicate the interpretation, reproducibility and comparison of findings from and across studies, particularly due to the lack of a consensus on the standardized and representative reference annotation. ZIKV reference genomes did not match virus strains sampled from the global epidemic or showed high level of heterogeneity in reported peptide lengths across their genome annotations.

To mitigate these concerns, we provided a correction with respect to the NCBI reference sequence NC_012532 for ZIKV (Figure S3). More information on the corrected reference sequence can be found at the Rega ZIKV reference sequence website ^5^.

**Figure S3:**
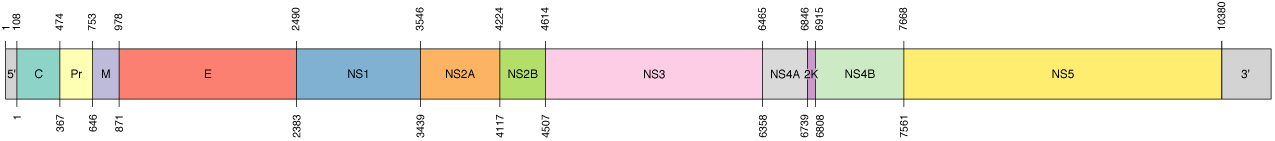
Accurate ZIKV genome positions of the different proteins based on corrections made in the the NCBI reference sequence NC_012532.

This example shows how the functionality of the XML configuration file, describing the genome annotation for all proteins, can greatly simplify the analysis to find associations of genomic features with clinically, epidemiologically or evolutionary parameters. We show that VIRULIGN makes this possible, while keeping manual processing limited to a minimum.

In particular, we replicate the evidence that indicated the necessity to correct the reference genome that was proposed by the NCBI. Extensive variation of a N-glycosylation motif in the Envelope (E) protein can be observed between different hosts, different viral lineages and even within a set of virus genomes derived from the historical MR766 strain.

##### S4.2.1 Sequence dataset

A specific subset of 19 (near-)complete ZIKV genomes was collected from Genbank, the resulting FASTA file ‘zikv.fasta’ can be found in the tutorial Zika folder:

https://github.com/rega-cev/virulign-tutorial/examples-alignments/ZIKV/

##### S4.2.2 Reference sequence

The reference genome sequence and the corresponding protein annotation for ZIKV can be found in the XML configuration file. This XML file ‘ZIKV-rega.xml’ can be found in the VIRULIGN references folder:

https://github.com/rega-cev/virulign/references

##### S4.2.3 Motif investigation

Previous literature has shown variability of the glycosylation motif around positions 150 165 in the E protein, and this has been suggested to result from excessive *in vitro* passaging [23]. We used VIRULIGN to create a position table of the amino acids in the alignment, annotated according to the respective protein. The following command was used:

~~~
virulign ZIKV-rega.xml zikv.fasta
   --exportKind PositionTable
   --exportReferenceSequence yes > zikv-aligned.csv
~~~

The resulting CSV file contains a column for each position in the genome, and can be found in the tutorial Zika folder:

https://github.com/rega-cev/virulign-tutorial/examples-alignments/ZIKV/

A simple R script shows the variability of the glycosylation motif across the different virus variants.

~~~
# import the alignment file
data<-read.csv(’zika-aligned.csv’,header=TRUE)
# determine the positions of the motif
reg<-match(’E_151’,names(data)):match(’E_163’,names(data))
# show the motif sequences
tidyr::unite(data[,c(1,reg)],Motif,-1,sep=’’)
~~~

The relevant region in the CSV file looks like:

~~~
id,E_151,E_152,E_153,E_154,E_155,E_156,E_157,E_158,E_159,E_160,E_161,E_162,E_163
Ref,M,I,V,N,D,T,G,H,E,T,D,E,N
KF268948,M,I,V,N,D,I,G,H,E,T,D,E,N
KF268949,M,I,V,N,-,-,-,-,-,-,D,E,N
KF268950,M,I,V,N,D,I,G,H,E,T,D,E,N
KU955595,M,I,V,N,D,T,G,H,E,T,D,E,N
KY014323,M,I,V,N,D,T,G,H,E,T,D,E,N
KU963574,M,I,V,N,-,-,-,-,-,-,D,E,N
HQ234500,M,I,V,N,-,-,-,-,-,-,D,E,N
KX369547,M,I,V,N,D,T,G,H,E,T,D,E,N
….
~~~

When additional meta-data is given as well, this analysis clearly illustrates the presence of a VNDT motif in viruses sampled from the recent epidemic, and an independence of the deletion regarding the host, year of collection and viral lineage (Figure S4).

**Figure S4:**
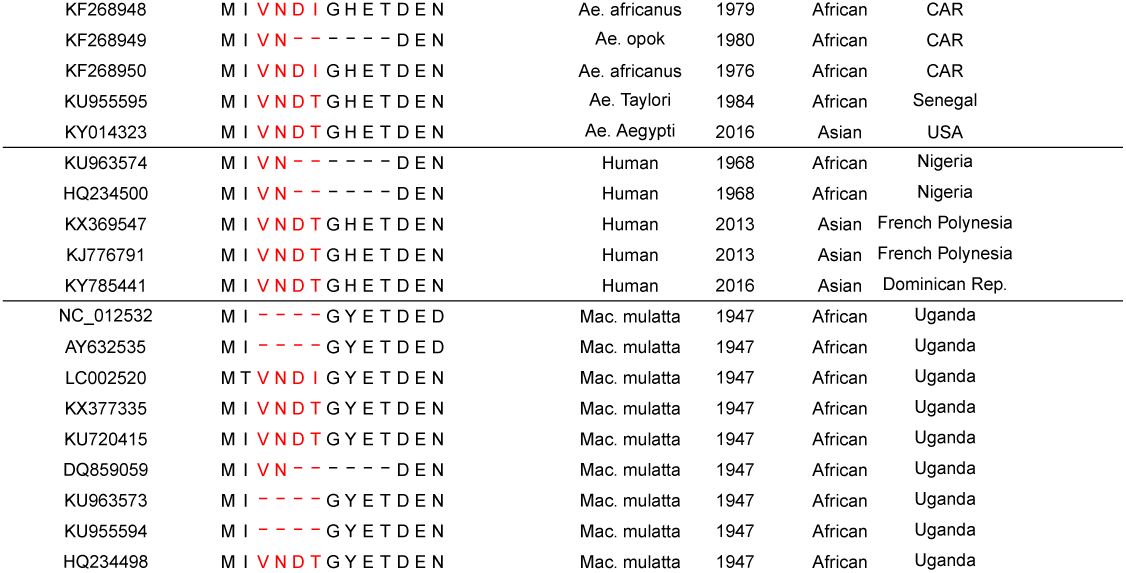
Variation in the glycosylation motif (red) of the ZIKV E protein across different hosts, virus lineage as well as date and country of collection.

##### S4.2.4 Scripts to select individual proteins

An alignment CSV table, that includes protein and position data in the header, can be easily processed by external tools. One could easily develop their own scripts to operate on this format, but many interesting manipulations can be done using default command line tools as well.

As an example, based on the whole-genome ZIKV alignment above, we can easily select a particular protein (e.g., the NS3 protein), using csvkit^6^.

~~~
# define index of the NS3 headers
ns3_headers=‘csvcut -n ZIKA-pos.csv | grep NS3 | cut -d “:” -f 2‘
# use comma as separator between header columns
ns3_headers=‘echo $ns3_headers | tr ’ ’ ’,’‘
# extract the NS3 region from the whole-genome alignment
csvcut -c “seqid,${ns3_headers}” ZIKA-pos.csv > ZIKA-NS3-pos.csv
~~~

#### S4.3 HIV-1 alignment: gag

The genome structure of HIV-1 is characterized by three ORFs, where each frame determines different genes encoding the viral proteins. The gag/pol/env gene organization, as for other retroviruses, encodes for important structural proteins and enzymes, which are first translated as large poly-proteins. Gag and pol have overlapping ORFs, requiring a ribosomal frame shift to reveal the pol ORF. HIV-1 gag encodes for several structural proteins and is considered as a potential target for antiretroviral treatment [21].

##### S4.3.1 Sequence dataset

HIV sequences were obtained from a large-scale analysis of HIV-1 diversity [10], where 2966 sequences from different HIV-1 subtypes have been analyzed. The FASTA input file ‘hiv.fasta’ can be found in the tutorial HIV-1 folder:

https://github.com/rega-cev/virulign-tutorial/examples-alignments/HIV-1/

##### S4.3.2 Reference sequence

The HXB2 sequence NC_001802^7^ was used as the reference genome. An XML file with the corresponding coding sequence and annotation is available for the different polyproteins of relevance. The respective files ‘HIV-HXB2-env.xml‘, ‘HIV-HXB2-gag.xml’ and ‘HIV-HXB2-pol.xml’ are available in the VIRULIGN references folder:

https://github.com/rega-cev/virulign/references

For this example, we used the file ‘HIV-HXB2-gag.xml‘.

##### S4.3.3 Alignment

To align the gag sequences, we used the following command

~~~
virulign HIV-HXB2-gag.xml HIV.fasta
    --exportKind GlobalAlignment
    --exportAlphabet Nucleotides
    --exportReferenceSequence yes > HIVgag.fasta
~~~

An option of VIRULIGN can be used to avoid insertions towards the reference sequence from being exported. This feature can be used to inspect the quality of the sequence dataset when insertions are sparse throughout the dataset or when insertions are not expected.

~~~
virulign HIV-HXB2-gag.xml HIV.fasta
  --exportKind GlobalAlignment
  --exportAlphabet Nucleotides
  --exportWithInsertions no
 --exportReferenceSequence yes > HIVgag-NoInsertions.fasta
~~~

#### S4.4 HIV-1 alignment: pol

A second example of HIV-1 alignment is directed towards drug resistance detection, which is still a major need for successful treatment, in particular in developing countries as a result of the up-scale of antiretroviral treatment. Therefore, the identification, understanding and interpretation of resistance mutations remains an important research topic. The pol polyprotein is cleaved into three viral enzymatic proteins (protease, reverse transcriptase, and integrase), each of which is an important drug target.

##### S4.4.1 Sequence dataset

We downloaded a large set of reverse transcriptase sequences (N=111223) from the Stanford University HIV Drug Resistance Database [17], heterogeneous in length and mapping of the complete reverse transcriptase region. The resulting FASTA file ‘HIVdb.fasta’ can be found in the tutorial HIV-1 folder:

https://github.com/rega-cev/virulign-tutorial/examples-alignments/HIV-1/

##### S4.4.2 Reference sequence

The HXB2 sequence NC_001802^8^ was used a reference genome. An XML file with the corresponding coding sequence and annotation is available for each specific ORF. The respective files ‘HIV-HXB2-env.xml‘, ‘HIV-HXB2-gag.xml’ and ‘HIV-HXB2-pol.xml’ are available in the VIRULIGN references folder:

https://github.com/rega-cev/virulign/references

For this example, we use the file ‘HIV-HXB2-pol.xml‘.

##### S4.4.3 Alignment

~~~
virulign HIV-HXB2-pol.xml HIVdb.fasta
  --exportKind GlobalAlignment
  --exportAlphabet Nucleotides
  --exportWithInsertions no
  --exportReferenceSequence yes > HIVrt.fasta
~~~

111184 sequences could be aligned by VIRULIGN, and we visually inspected the quality of the alignment. The constructed alignment can be used as input for different applications to investigate drug resistance mutations identification and interpretation. Thanks to VIRULIGN’s computional complexity, our new method is able to deal well with large dataset what is reflected in favorable run-times for this particular analysis. VIRULIGN performed this alignment in 49 minutes, while it took MAFFT 10 hours and 50 minutes9 10.

#### S4.5 Computational complexity

We derive the computational complexity of VIRULIGN by inspecting the algorithm described in the methods section of the main manuscript. We observe that for each target sequence *t* ∈ 𝒯, a constant number of Needleman-Wunsch alignments is performed. It is well known that the computational complexity of a Needleman-Wunsch alignment of a sequence tuple (*s*_1_, *s*_2_) is 𝒪(|*s*_1_| · |*s*_2_|) in both space and time [13]. As in VIRULIGN, we consider a reference sequence *r* and a set of target sequences 𝒯, and each target sequence *t* is aligned to *r*, the maximal Needleman-Wunsch computational complexity 𝒪(*NW*_*max*_) is thus 𝒪(|*r*| · *max*({|*t*| | *t* ∈ 𝒯 })). As this applies to all target sequences, VIRULIGN’s complete computation complexity is | 𝒯 | · 𝒪(*NW*_*max*_), or simply 𝒪(| 𝒯 | · |*r*| · *max*({|*t*| | *t* ∈ 𝒯 })).

#### S4.6 List of case study examples

VIRULIGN has been used for a large number of analyses with respect to virus genomics. We provide a non-exhaustive list of examples:

- Identification of HIV-1 drug resistance mutations and the pathways emerging under drug selective pressure, as well as modeling **the different factors leading to treatment failure and trends over time** [14,24,27,26,4,5].
- Large-scale analysis of HIV-1 and HCV sequence datasets to explore genetic diversity at population level and to map structural and functional factors that shape viral evolution [1,10,2,3].
- Evaluation of the annotation and representativeness of current reference genomes, and support of correcting the NCBI reference sequence for the Zika virus [23].
- Web-application to support surveillance and tracing of viral outbreaks (e.g., HIV-1 and DENV), necessitating efficient analysis of large sequence databases and phylogenetic trees [12].
- VIRULIGN was recently integrated into the RegaDB data management and analysis platform for the clinical follow-up of HIV-1 patients [27,11].
- Evaluation of an automated framework for the virus typing (HIV-1, Dengue, Zika and other viruses) and resistance interpretation algorithms [15,20,22]

## Footnotes

1 To print these options, invoke the virulign command without any arguments.

2 www.ncbi.nlm.nih.gov/genomes/VirusVariation

3 www.ncbi.nlm.nih.gov/nuccore/NC_001477

4 Performed on a 3.6 GHz Intel^®^ Core i7 CPU with 12 GB of RAM, where each application had access to 1 CPU core.

5 rega.kuleuven.be/cev/reference-sequences/rega-zikv

6 https://github.com/wireservice/csvkit

7 www.ncbi.nlm.nih.gov/nuccore/NC_001802

8 www.ncbi.nlm.nih.gov/nuccore/NC_001802

9 Clustal Omega and MUSCLE did not finish in more than 5 days.

10 Performed on the same hardware configuration as declared earlier.

